# Effective connectivity reveals dual-route mechanism of visual prediction precision via insula and pulvinar

**DOI:** 10.1101/2025.07.07.662254

**Authors:** Linzhi Tao, Trevor Steward, Joshua Corbett, Rebecca K. Glarin, Tudor V. Sava, Marta I. Garrido

## Abstract

The brain’s ability to weight predictions by their precision is a central mechanism in predictive processing, enabling optimal integration of prior expectations with incoming sensory input. Despite its theoretical significance, the neural circuitry that implements precision-weighted prediction remains unclear. Using 7-Tesla fMRI and dynamic causal modelling (DCM), this study investigated how the brain encodes the precision of predictions during a visual cueing task with high- and low-precision conditions. We focused on the key regions implicated in predictive processing: the insular cortex, the pulvinar nucleus of the thalamus, and primary visual cortex (V1). Behaviourally, participants showed significantly greater accuracy in the high-precision condition (*p* < .001), confirming effective task manipulation. DCM analyses revealed that high-precision predictions elicited excitatory modulation of connectivity from the insula to V1 (Pp = .95), alongside inhibitory influences from the insula to the pulvinar (Pp = .99) and from the pulvinar to V1 (Pp = .89). Furthermore, leave-one-out cross validation revealed that individual differences in behavioural sensitivity to precision were positively predicted by pulvinar-to-insula connectivity (*r* = .36, *p* = .026) and negatively predicted by the connectivity between pulvinar and V1 (pulvinar to V1: *r* = .35, *p* = .033; V1 to pulvinar: *r* = .37, *p* = .026), highlighting the behavioural relevance of these pathways. Together, these findings suggest a dual-route mechanism whereby the insula directly enhances top-down predictions in V1 while indirectly dampening bottom-up sensory input via the pulvinar. This mechanism may facilitate Bayesian integration under uncertainty and offers new hypotheses into how precision weighting may be disrupted in neuropsychiatric conditions.

## Introduction

The human brain continuously engages in predictive processing, a fundamental mechanism through which it anticipates sensory inputs and refines perception via feedforward and feedback signals (Friston, 2005). Central to this framework is the notion of hierarchical Bayesian inference, wherein the brain constructs models of the environment, updates internal models based on sensory evidence, and weighs the reliability of incoming information by estimating its precision (the inverse of variance; Clark, 2013; K. Friston, 2012). This dynamic precision-weighting mechanism is critical for perception as it determines whether the brain prioritises sensory input or top-down predictions. Dysregulation in precision modulation has been proposed as a key mechanism in neuropsychiatric disorders, such as autism and schizophrenia (Haarsma et al., 2021; Lawson et al., 2014; Pellicano & Burr, 2012). However, the underlying neural circuitry that enables precision-related computations remains unknown. In particular, it is unclear how different brain regions dynamically interact to shape prediction and its associated precision.

Three key regions implicated in feedforward and feedback signalling in visual predictive processing are the primary visual cortex (V1), the insula, and the pulvinar. While each of these regions has been associated with distinct aspects of prediction (Ficco et al., 2021; Kanai et al., 2015; Siman-Tov et al., 2019), their dynamic interactions in precision modulation remain underexplored. V1, the earliest cortical recipient of visual input, integrates bottom-up feedforward signals from the retina and top-down feedback signals from higher-order regions (Petro et al., 2014). These top-down feedback signals have been proposed to carry precision-related information, modulating V1’s response to sensory input (Tani & Kashimori, 2021). This hypothesis was further supported by a laminar fMRI study showing that prior expectations evoked activity in the deep layers of V1 (Aitken et al., 2020; Kok et al., 2016), which typically receive top-down feedback signalling (Briggs, 2020), implicating top-down influences from higher regions in the processing of prior expectations. However, the source region of this expectation-related feedback signalling remains unidentified. Moreover, it is unclear whether this feedback specifically relates to precision modulation.

One potential source of the precision-related feedback to V1 is the insula. A common theme across the multiple functional roles of the insula is integrating both the salience and relevance of the stimuli and modulating sensory processing accordingly, where highly salient information can be prioritised for processing (Menon & Uddin, 2010; Seth, 2013). In the context of visual prediction, the insula may exert top-down modulation on early visual areas to prioritise high-precision information. In support of this, a 7-Tesla (7T) fMRI study on interoception showed reduced insula activity to visual stimuli that were expected by the participants (Salomon et al., 2016), potentially reflecting top-down inhibition of feedforward signals to promote internally generated, high-precision predictions. This cortico-cortical modulation would allow the brain to enhance or suppress visual predictions dynamically, optimising perceptual accuracy under varying levels of prediction precision.

The pulvinar, a collection of higher-order thalamic nuclei, has also been proposed to regulate visual processing in V1 for precision modulation (Kanai et al., 2015). Anatomically, the pulvinar has extensive reciprocal connections with V1, which supports its ‘modulator’ role in regulating V1 activity (Cortes et al., 2024). Precision weighting may occur through feedback projections from the pulvinar to V1. An electrophysiological study further demonstrated that inactivating the pulvinar nuclei reduced responsiveness in the superficial layers of V1 to visual stimulation (Purushothaman et al., 2012). Since the superficial layers are the primary source of feedforward projections to higher cortical areas (Yang et al., 2021), this finding suggests that the pulvinar may gate V1’s feedforward signalling to higher cortical areas, possibly regulating the propagation of precision-weighted predictions. Nevertheless, whether this gating function specifically relates to precision modulation requires further examination.

A central challenge, therefore, is to identify how the insula, pulvinar, and V1 interact during visual predictive processing under varying levels of precision. To this end, dynamic causal modelling (DCM; Friston et al., 2003) offers a powerful framework for investigating the effective connectivity between brain regions (i.e., the directed influence of one brain region on another). Crucially, understanding the directionality of these influences helps disentangle whether changes in neural activity reflect feedforward or feedback signalling, thereby providing mechanistic insights into precision modulation in the brain.

This study employed DCM on 7T fMRI data acquired during a visual cueing task with high- and low-precision conditions to investigate the modulatory connections between V1, the insula, and the pulvinar. Behaviourally, we hypothesised that participants’ response accuracy would be higher in the high-precision condition than in the low-precision condition. We tested a DCM model that encompassed the following hypotheses motivated by the literature reviewed above: (1) the insula exerts inhibitory modulation on V1 during high-precision predictions; (2) the pulvinar exerts inhibitory modulation on V1 during high-precision predictions; and (3) the strength of these modulations predicts inter-individual variability in behavioural performance.

## Methods

### Participants

Thirty-one healthy individuals with normal or corrected-to-normal vision participated in the experiment. All participants met the following eligibility criteria: (i) aged between 18 and 50 years, (ii) were fluent English speakers, and (iii) had no contraindication to MRI (e.g., metal implants, claustrophobia, pregnancy). Participants attended a single session at the Melbourne Brain Centre Imaging Unit (The University of Melbourne, Parkville). Of the original sample, all data from one participant and partial data from three participants (which included two runs from one participant’s data, and one run from two participants’ data) were excluded due to excessive head movement during scanning (please see *Image Preprocessing* in the Methods section for more information). All data from one participant and partial data from two participants (which included two runs from two participants separately) were excluded due to poor behavioural performance (please see *Task Performance* in the Results section). The final sample consisted of 29 participants (16 female, 1 prefer not to say, mean age 27.8 ± 6.9 years). This study was approved by the University of Melbourne Human Research Ethics Committee (Ethics ID: 24710, reference number: 2022-24710-35008-4). All participants provided written informed consent and received monetary compensation for participation.

### MRI task design

We designed a visual probabilistic cueing paradigm using gratings (Figure 1) to elicit visual predictions and manipulate their precision. At the start of each trial, participants were presented with a coloured cue (blue, yellow, or green) at the centre of the screen for 750 ms, followed by a blank fixation circle for 500 ms. The first grating was then presented for 250 ms, followed by another blank fixation circle for 500 ms, and then the second grating for 250 ms. This was followed by a blank fixation circle for 2500 ms, and then an instruction to respond to whether the two gratings are identical or different. Participants responded using an MRI-compatible button box placed in their right hand.

**Figure 1.**
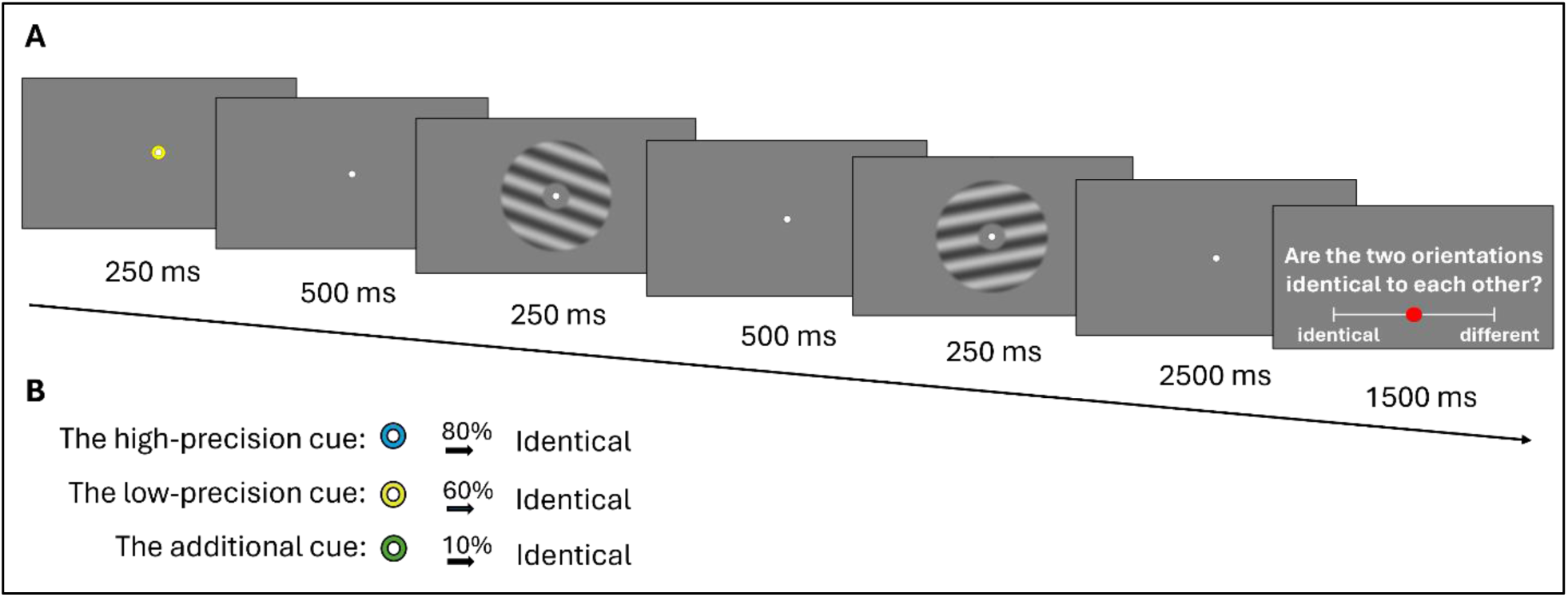
A visual probabilistic cueing paradigm using gratings. (A) Participants were first presented with a visual cue with varying prediction precision, followed by two consecutive gratings with either identical or non-identical orientations. Participants were then asked to respond whether the two stimuli were the same or different using a two-key button box. (B) There were three cues predicting identical orientations with different precision. The high-precision cue predicts identical orientations 80% of the time. The low-precision cue predicts identical orientations 60% of the time. The additional cue was used to balance the overall expectation of seeing identical gratings. As the number of trials was the same for each cue, the probability of seeing identical orientations over all trials was (80% + 60% + 10%)/3 = 50%. Participants were also told prior to scanning that, overall, the gratings were identical half of the time and different half of the time.

Each cue predicted whether the two gratings were identical with different levels of precision: (1) the *high-precision cue* predicted identical gratings in 80% of the trials; (2) the *low-precision cue* predicted identical gratings in 60% of the trials; and (3) an *additional cue* predicted identical gratings in 10% of the trials. The 10% precision cue was used to balance the overall expectation of seeing identical gratings. By including the 10% cue, and ensuring that all cues occurred equally frequently, the overall probability of identical gratings across all trials was reduced to 50% (i.e., (80% + 60% + 10%)/3). Participants were also informed that, on average, the gratings would be identical in half of the trials and different in the other half.

To encourage reliance on both the cue-based predictions and sensory evidence from the two gratings presented, we deliberately made the difference between the two gratings subtle in most trials (1 to 8 degrees). This ensured that visual information alone was often insufficient to determine whether the gratings were identical, requiring participants to integrate prior knowledge from the cues with the visual input to make their judgments.

There were 360 trials in total, 120 trials for each cue condition. This was divided into three runs, with 12 blocks in each run, each containing 10 trials. Different cue conditions were randomised across the entire experiment. The sequence of the conditions was optimised using the Neurodesign algorithm (Durnez et al., 2018). The inter-stimulus interval (ISI) was jittered around a mean of 2.5 s. The average length of a trial was 6.25 s. The break between blocks was 5 s, where the feedback of the performance in the previous block was presented (i.e., the number of correct responses in the previous ten trials). The break between runs was around 2 min. The total length of the scanning session was approximately 50 min.

For each condition, each participant’s response accuracy was calculated by dividing the number of correct responses by the total number of responses in that condition. A paired-samples t-test was then conducted to compare response accuracy between the high- and low-precision conditions, assessing whether the task elicited a significant behavioural difference between the two conditions.

### Image acquisition

A Siemens Magnetom 7T Plus MRI Scanner (Siemens Healthcare, Erlangen, Germany) equipped with a 32Rx/1Tx channel head coil (Nova Medical Inc., Wilmington MA, USA) was used. The functional imaging (T2*-weighted) was an accelerated GRE-EPI sequence in the steady state (Moeller et al., 2010; 84 interleaved slices parallel to the anterior-posterior commissure line, slice thickness = 1.6 mm (no spacing between slices), TR = 0.8 s, TE = 22.2 ms, flip angle = 45°, field of view (FOV) = 20.8 cm, matrix size = 130 × 130, volume = 1084), anterior to posterior phase direction. The acceleration parameters consisted of a multiband factor of 6 and a Generalised Autocalibrating Partially Parallel Acquisitions (GRAPPA) factor of 2 (Griswold et al., 2002). For each participant, a high-resolution structural image (T1-weighted) was acquired using Magnetization Prepared 2 Rapid Acquisition Gradient Echoes sequence (MP2RAGE; Marques et al., 2010) to assist with functional image co-registration and normalisation (224 sagittal slices aligned parallel to the midline, slice thickness = 0.75 mm (no spacing between slices), TR = 5 s, TE = 2.04 ms, TI = 700/2700 ms, flip angle = 4°,5°, FOV = 24 cm, matrix size = 320 × 320). Standard foam pads were placed at both sides of the participants’ head to minimise head movement during the scan. The task was presented using an MRI-compatible screen. Pulse and respiratory data were collected during each functional sequence with Siemens Bluetooth equipment.

### Image preprocessing

MRI data were preprocessed with Statistical Parametric Mapping (SPM) 12 (Wellcome Trust Centre for Neuroimaging, London, UK) in MATLAB 2023a (The MathWorks Inc., Natick, MA) on the high-performance computing platform (HPC), Spartan, at the University of Melbourne. Motion artifacts were corrected by realigning individual’s time series to their mean image using least-squares minimisation and a six-parameter rigid-body spatial transformation. The 4th Degree B-Spline interpolation method was used in image resampling. The Motion Fingerprint toolbox (Wilke, 2012) was used to calculate the total displacement (TD; a single measure that takes into consideration both the translations and rotations) of participants’ head movement. Of the original sample, all data from one participant and partial data from three participants (which included two runs of one participant’s data, and one run of two participants’ data) were excluded due to excessive head movement (mean TD > 2 mm). Physiological noise was corrected by including the regressors for respiration and pulse created by the PhysIO Toolbox (Kasper et al., 2017) in the first-level GLM models. For each participant, the anatomical images were coregistered to the corresponding mean functional image. Segmentation and normalisation were then applied using the International Consortium of Brain Mapping template (European brain) with the DARTEL approach in SPM (Ashburner, 2007). The resulting normalised anatomical volumes were then used to normalise the functional images to the Montreal Neurological Institute (MNI) space for each participant.

### Dynamic causal modelling

#### Overview

Dynamic causal modelling (DCM; Friston et al., 2003) is a forward modelling technique that can be used to infer endogenous connections and task-dependent modulatory connections between brain regions, that is, the directed causal influence of one region on other regions. For fMRI studies, the model uses three sources of variance (i.e., neural activities, hemodynamic changes, and observation noise for fMRI data) to predict the observed fMRI timeseries (Zeidman, Jafarian, Corbin, et al., 2019). The neural component of the model specifies two connectivity matrices (i.e., the endogenous connections and task-related modulatory connections between regions), which represents the belief about the underlying neural activity between the regions that explains the variance in the fMRI timeseries (Friston et al., 2003; Zeidman, Jafarian, Corbin, et al., 2019). Models with different combinations of the connectivity parameters can be constructed based on specific research questions and hypotheses. The models are then estimated through Bayesian model inversion that outputs the estimated strength of the connectivity and their posterior probability (Pp; a measure that indicates how likely the strength of a parameter is different from zero).

#### First- and second-level GLM for DCM analysis

A general linear model (GLM) was specified for the DCM analysis concatenating the three task runs into a single run using the spm_fmri_concatenate() function. The baseline regressors for each of the three runs were included in the design matrix.

The first column of the matrix specifies the onsets of all trials (including all precision conditions), which was later used as the driving input to the DCM model. This condition is referred to as the TASK condition. The high- and low-precision conditions were modelled as separate regressors in the design matrix, where the high-precision condition was later used as the modulatory input to the DCM model. The contrast between the high- and low-precision conditions at the second-level GLM was used to identify the activations related to the difference in precision. The design matrix also included regressors for the instruction period, the response period, and the feedback period, as well as regressors for physiological noise corrections and motion correction.

For each participant, the onset and duration of each trial were convolved with a canonical hemodynamic response function to create regressors for the TASK, high-precision and low-precision conditions in the first-level GLM. Trial onsets were defined as the onset of the cue presentation. Trial durations spanned from the onset of the cue to the offset of the second grating. This time window was selected to isolate neural activity related to the predictive process (encompassing both the cue and grating presentations) while minimising motion-related artefacts from the finger-pressing response. A high-pass filter at 128 s was used to remove low-frequency noise. Local temporal autocorrection (FAST; Corbin et al., 2018) was applied.

Parameter estimates of the GLM were computed for each voxel for the effect of TASK, which includes all precision conditions, and for the contrast between the high- and low-precisions. An F-contrast of the effect of interest of the task was estimated and later used in timeseries extraction to regress out the nuisance effects that are not of interest in the DCM analysis.

A second-level GLM analysis was conducted to test if BOLD responses are significantly different between the high- and low-precision conditions across participants. A significance thresholding of whole-brain FDR (Benjamini & Hochberg, 1995) < .05 was applied to correct for multiple comparisons (Lindquist & Mejia, 2015).

#### Region of interest selection and time-series extraction

Regions of interest (ROIs) were defined by a conjunction map of the anatomical mask of the hypothesised regions and the activation map of the contrast between the high- and low-precision conditions (high > low) in the GLM model. The anatomical masks were created using the JuBrain Anatomy toolbox in SPM12 (Eickhoff et al., 2005). A representative timeseries was extracted by computing the principal eigenvariate of the activated voxels enclosed by the anatomical mask for each ROI for each participant. The timeseries were pre-whitened, high-pass filtered, and the nuisance effects not covered by the F-contrast of interest were regressed out of the timeseries, following published DCM processing guidelines (Zeidman, Jafarian, Corbin, et al., 2019).

#### Model specification, estimation, and selection

A full model of all the possible endogenous and modulatory connections between regions were specified in the DCM module in SPM12 (Figure 4B). This includes an A-matrix specifying all the possible endogenous connections between regions, as well as the self-connections of all regions. The high-precision condition was used as the modulatory input to all possible connections between regions in the model (specified in the B-matrix). The TASK condition was used as the driving input to the model (specified in the C-matrix). Both the V1 and pulvinar were switched on in the C-matrix in the full model as both regions would be reasonable candidates receiving the visual driving input being early regions in the visual processing hierarchy (Purushothaman et al., 2012).

The full DCM model was specified and estimated for each participant. The first-level modulation parameter estimates were then modelled and estimated at the second level using parametric empirical bayes (PEB; Zeidman, Jafarian, Seghier, et al., 2019), which calculates the probability of the model parameters given the observed fMRI timeseries that accounts for both the estimated strength of the connectivity and the estimated uncertainty of the connectivity strengths at the first level.

Bayesian model selection (BMS; Stephan et al., 2009) was used to test whether driving inputs to V1, the pulvinar, or both, showed stronger model evidence. Three families of models were specified and compared in BMS: (1) both V1 and pulvinar, (2) V1 only, and (3) pulvinar only. The winning family was carried forward for subsequent analysis.

Bayesian model reduction (BMR; Friston et al., 2016) was then applied to evaluate all reduced models of the winning model of the BMS. Parameters of the connectivity that did not contribute sufficiently to the model evidence were pruned away during the iterative process of BMR over all the possible reduced models (Friston et al., 2016; Zeidman et al., 2019). Bayesian model averaging (BMA; Friston et al., 2016; Zeidman, Jafarian, Seghier, et al., 2019) was computed over modulation parameters of the reduced models, where their relative weighting in the averaging process was adjusted by their model evidence. The BMA results were thresholded to include parameters that showed ‘positive evidence’ (posterior probability > .75; as labelled by Kass and Raftery, 1995) of being different from zero at the second level.

#### Predicting the high-precision effect using strength of the modulatory connections

The high-precision effect (i.e., the accuracy gain of high precision) was computed by taking the difference between participants’ accuracy in the high-precision condition and the mean accuracy throughout the task. This behavioural measure indicates the extent to which a participant performed better in the high-precision condition than their average across conditions. This measure was included in the PEB model as a behavioural covariate to test if changes in the high-precision effect were explained specifically by changes in the strength of any of the modulations (elicited by high precision), linking the observed behavioural and neural effects of precision. A positive parameter estimate indicates that participants who showed stronger modulation (between the modelled regions) elicited by high precision would also tend to show larger high-precision effect.

Modulation parameters with positive evidence (Pp > .75) were further examined in leave-one-out cross validation to test if the effect size of the modulations would be sufficiently large to predict participants’ high-precision effect (Zeidman et al., 2019), and if the effect would be robust against outliers. At every iteration of the cross validation, a PEB model was estimated while leaving out a participant and was used to predict the behavioural covariate (Zeidman et al., 2019). The predicted value of the behavioural covariate included in the PEB model was estimated for each left-out subject. The predicted scores of the covariate were then correlated with actual scores of the covariate to test if effect sustains throughout the leave-one-out iterations. A significant correlation (*p* <.05) between the predicted and actual scores indicates that the effect size of the modulatory connections is sufficiently large to predict the behavioural covariate.

## Results

### The task elicited behavioural difference between the high- and low-precision conditions

Out of the original sample (*n* = 31), one participant performed below 50% accuracy in all runs of their scans, and two participants performed below 50% accuracy in two of their scans.

Runs with accuracy below 50% were excluded from the analyses. Together with the exclusion of one participant due to excessive motion during the scan, we included a sample of 29 participants, with a mean response accuracy of 66.1% (SD = 6.8%). The mean reaction time was 0.34 s (SD = 0.18 s).

A Shapiro-Wilk test indicated that the difference scores in participants’ response accuracy between the high- and low-precision conditions were normally distributed (*W* = .97, *p* = .960). As expected, participants showed significantly higher response accuracy in the high-precision condition compared to the low-precision condition, *t*(53.2) = 5.61, *p* < .001 (Figure 2).

**Figure 2.**
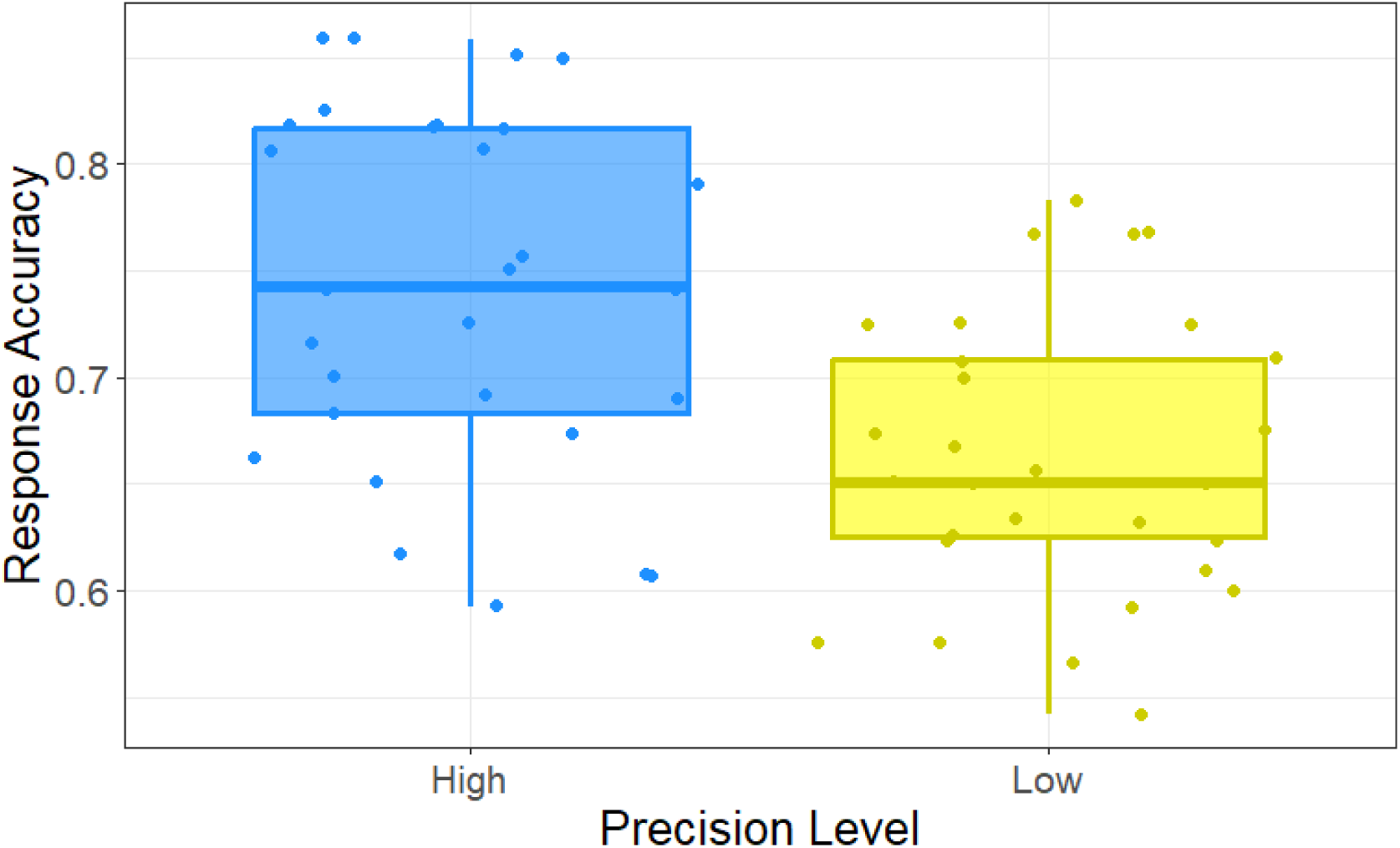
Response accuracy of the high- and low-precision conditions. Participants’ response accuracy was significantly higher in the high-precision condition than the low-precision condition, *t*(53.2) = 5.61, *p* < .001. Each dot represents a participant’s response accuracy.

### Second-level GLM: Differences in V1, pulvinar and insula activation between high- and low-precision conditions

The second-level GLM results of the contrast between the high- and low-precision conditions (high>low) showed activations (thresholded at *p* < .05, pTFCE-corrected; Spisák et al., 2019) in the hypothesised regions (V1, pulvinar, insula) as well as a number of other regions including primarily the superior medial frontal gyrus, the superior occipital gyrus, cerebellum lobule VI, and the ventral lateral nucleus in the thalamus (Figure 3 and Table S1). These activations did not survive the whole-brain FDR correction (*p* < .05, FDR).

**Figure 3.**
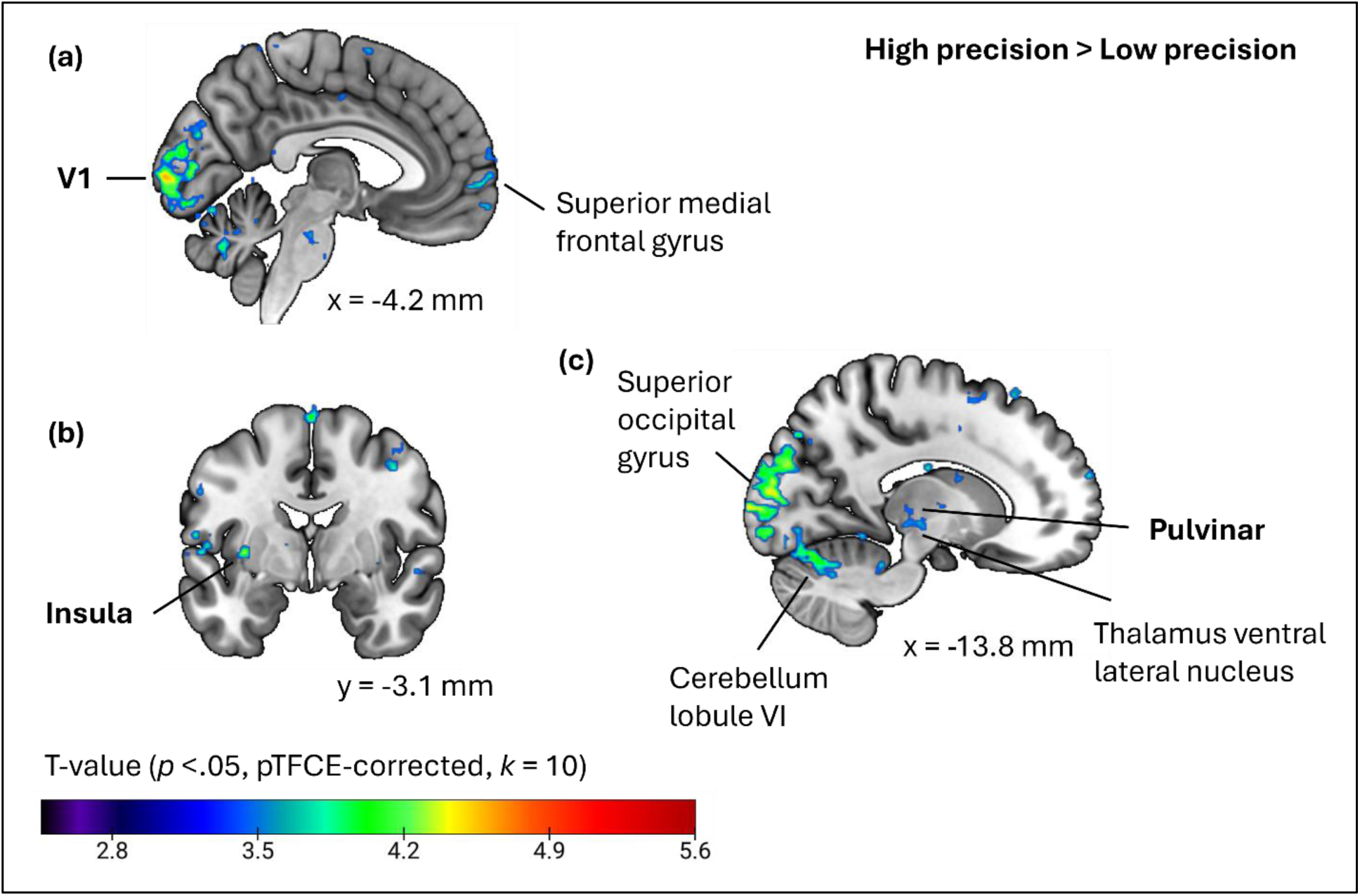
Second-level whole-brain statistical map of the contrast between the high- and low-precision conditions (High > Low). Activations were thresholded at *p* < .05, pTFCE-corrected (Spisák et al., 2019), with a minimum cluster size (k) = 10. The colour bar represents corresponding t-values. Numbers represent Montreal Neurological Institute (MNI) coordinates. From (a) to (c), the numbers indicate the x-axis coordinate (x = −4.2 mm), the y-axis coordinate (y = −3.1 mm), and the x-axis coordinate (x = −13.8 mm), respectively. V1: primary visual cortex.

### Regions of interest and the full model

Three regions of interest (ROIs) were defined by a conjunction map between the anatomical mask of the hypothesised regions and the activation map of the contrast between the high- and low-precision conditions (High > Low) in the second-level GLM analysis. To visualise the extent to which the ROIs overlap across participants, individual ROI masks were binarised and overlayed across participants (Figure 4A). Brighter colours indicate greater overlaps between participants. Figure 4B illustrates the full DCM model constructed from the three ROIs. In this model, the high-precision condition was used as the modulatory input to all possible connections between regions in the model. The TASK condition was used as the driving input to both V1 and the pulvinar in the full model.

**Figure 4.**
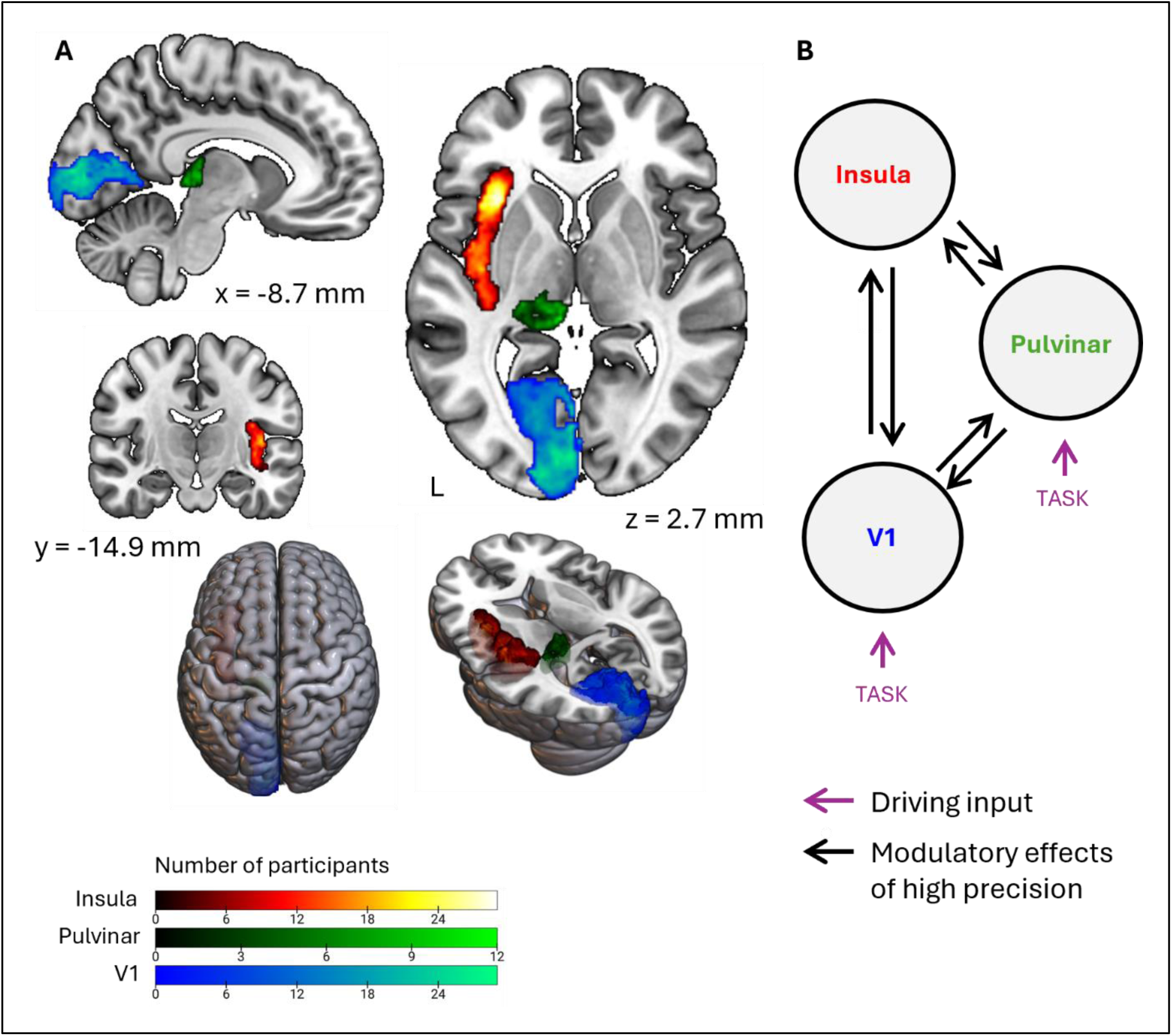
The regions of interest (ROIs) and the full dynamic causal modelling (DCM) model space. (A) The ROIs: the insula, pulvinar, and the primary visual cortex (V1). Left = Left (L). Numbers represent Montreal Neurological Institute (MNI) coordinates. For the sagittal view, the number indicates the x-axis coordinate (x = −8.7 mm). For the coronal view, the number indicates the y-axis coordinate (y = −14.9 mm). For the axial view, the number indicates the z-axis coordinate (z = 2.7 mm). The intensity of the masks indicates to what extent the individual ROI masks overlap with each other. This was created by overlaying the binarized ROI masks of each participant (*N* = 29). Lighter colours indicate greater overlaps between participants (i.e., more participants having a particular voxel included in their ROI mask). The individual ROIs were defined by a conjunction map of the anatomical mask of the hypothesised regions and the effect of interest map in the second-level GLM model. (B) The full DCM model. The high-precision condition was used as the modulatory input to the model. All possible connections between the three regions were switched on in the full model for modulatory effects. TASK (i.e., all the precision conditions) was used as the driving input to the model and applied to both the V1 and the pulvinar in the full model. Note that all the possible endogenous connections (including both the between-region connections and self-connections) were switched on in the full model but are not shown on the figure for simplicity.

### Bayesian model selection suggested driving input to V1

Three models were compared in BMS to infer whether the driving input should be applied to the V1 and/or pulvinar (Figure 5A): Model 1: driving input applied to both V1 and pulvinar; Model 2: driving input applied to V1; and Model 3: driving input applied to pulvinar. The BMS result (Figure 5B) showed that Model 2 (Pp = .56) outperformed Model 1 (Pp = .06) and Model 3 (Pp = .38), suggesting driving input to V1.

**Figure 5.**
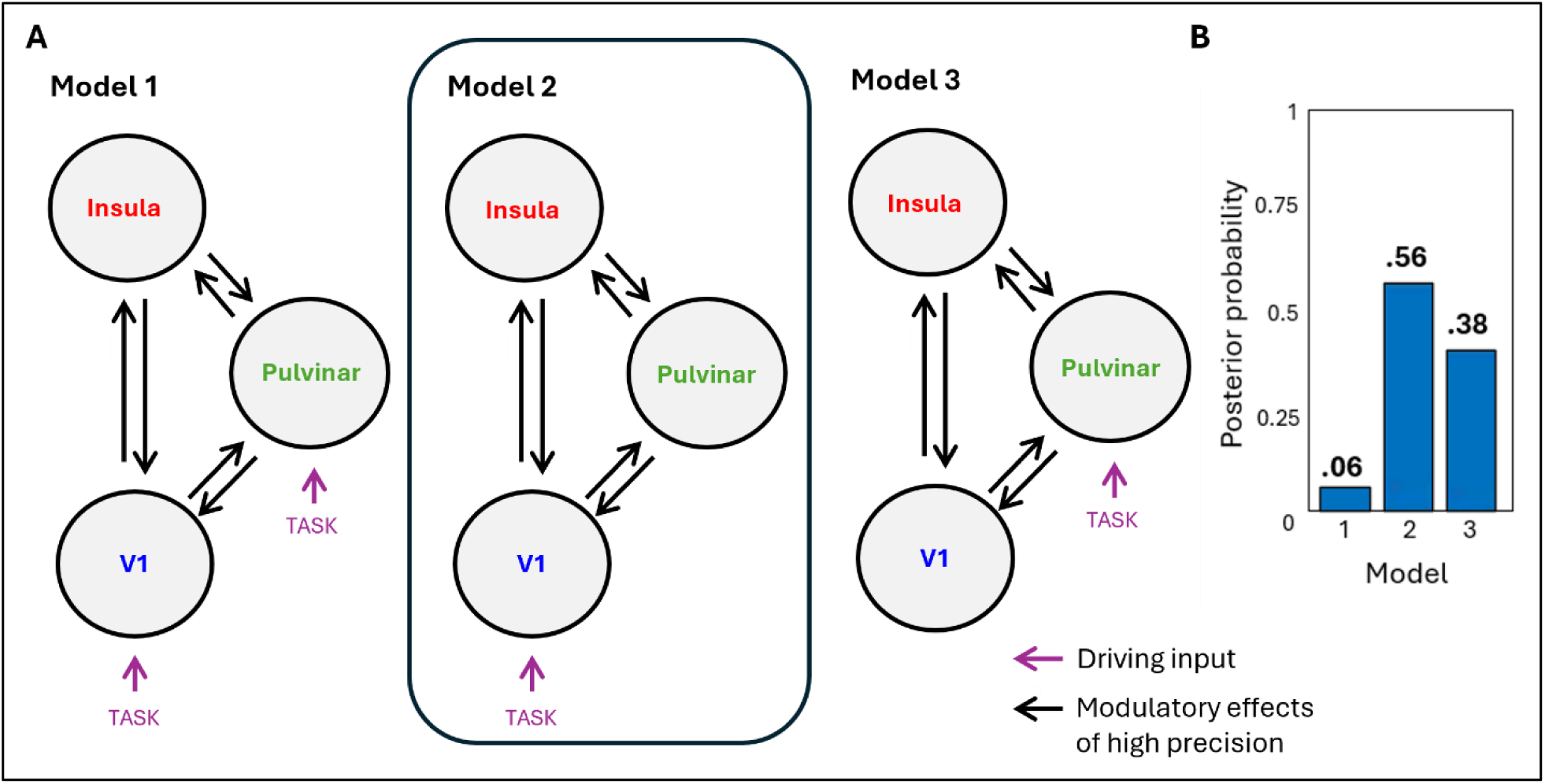
(A) Bayesian model selection (BMS) of three models: Model 1: driving input applied to both V1 and pulvinar; Model 2: driving input applied to V1; and Model 3: driving input applied to pulvinar. The endogenous and modulatory connections were all switched on for the three models. (B) Posterior probability of the three models computed in the BMS analysis reveal that the best model has the driving input into V1 (Model 2).

### PEB results revealed that high precision elicited modulatory connections across all three regions

The BMA results (Figure 6) showed three modulatory connections with posterior probability > .75, indicating “positive evidence” that the strength of the modulation is different from zero (Kass & Raftery, 1995). It was found that high precision predictions elicited (1) an excitatory modulation from the insula to V1 (Pp = .95), (2) an inhibitory modulation from the insula to pulvinar (Pp = .99), and (3) an inhibitory modulation from the pulvinar to V1 (Pp = .89). See Table 1 for a summary of the results for the modulatory connections.

**Figure 6.**
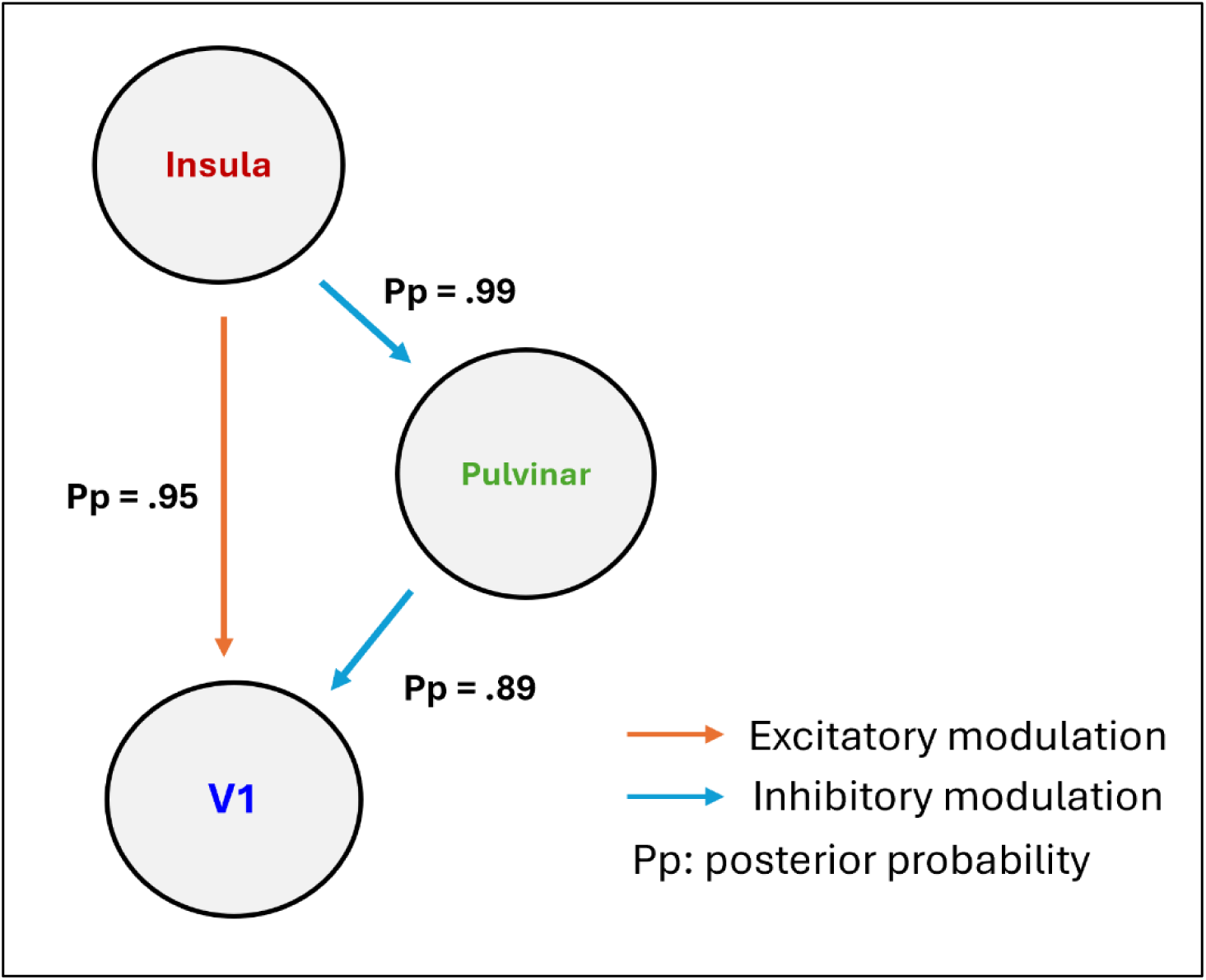
Modulatory effect of high precision. Arrows represent modulatory connectivity with > .75 posterior probability (Pp; the probability that the connectivity parameter is different from zero). V1: Primary visual cortex.

**Table 1.** Bayesian model averaging (BMA) results: DCM parameters for endogenous and modulatory connections. *Posterior probability (Pp) exceeding .75 indicates positive evidence. V1: primary visual cortex. Cp: posterior covariance. Ep: posterior expectation.

| Connection | Ep | Cp | Pp |
| --- | --- | --- | --- |
| **Endogenous connections (A-matrix)** |  |  |  |
| V1 → V1 | 0.14 | 0.0027 | .97* |
| V1 → Insula | 0.07 | 0.0021 | .84* |
| Pulvinar → V1 | 0.33 | 0.0065 | 1.00* |
| Pulvinar → Pulvinar | -0.58 | 0.0024 | 1.00* |
| Pulvinar → Insula | -0.17 | 0.0049 | .97* |
| Insula → V1 | 0.15 | 0.0028 | .98* |
| Insula → Pulvinar | 0.10 | 0.0008 | 1.00* |
| Insula → Insula | -0.23 | 0.0028 | 1.00* |
| **Modulatory connections (B-matrix) effect of high precision** |  |  |  |
| Pulvinar → V1 | -0.37 | 0.0682 | .89* |
| Insula → V1 | 0.43 | 0.0394 | .95* |
| Insula → Pulvinar | -0.52 | 0.0225 | .99* |
| **Effect on predicting the high-precision effect** |  |  |  |
| V1 → Pulvinar | -4.23 | 2.6756 | 0.94* |
| Pulvinar → V1 | -14.32 | 9.067 | 1.00* |
| Pulvinar → Insula | 3.61 | 12.3271 | 0.82* |
| Insula → Pulvinar | 8.24 | 6.2385 | 0.98* |

### The strength of the modulatory connections predicted the high-precision effect

The behavioural covariate, high-precision effect, was included in the model to further investigate if the strength of the modulatory connections could predict the magnitude of the behavioural effect of high precision. This covariate was computed as the difference of a participant’s response accuracy between high-precision condition and their average response accuracy. The BMA results showed that four modulatory connections predicted the high-precision effect with above .75 posterior probability (Figure 7A).

**Figure 7.**
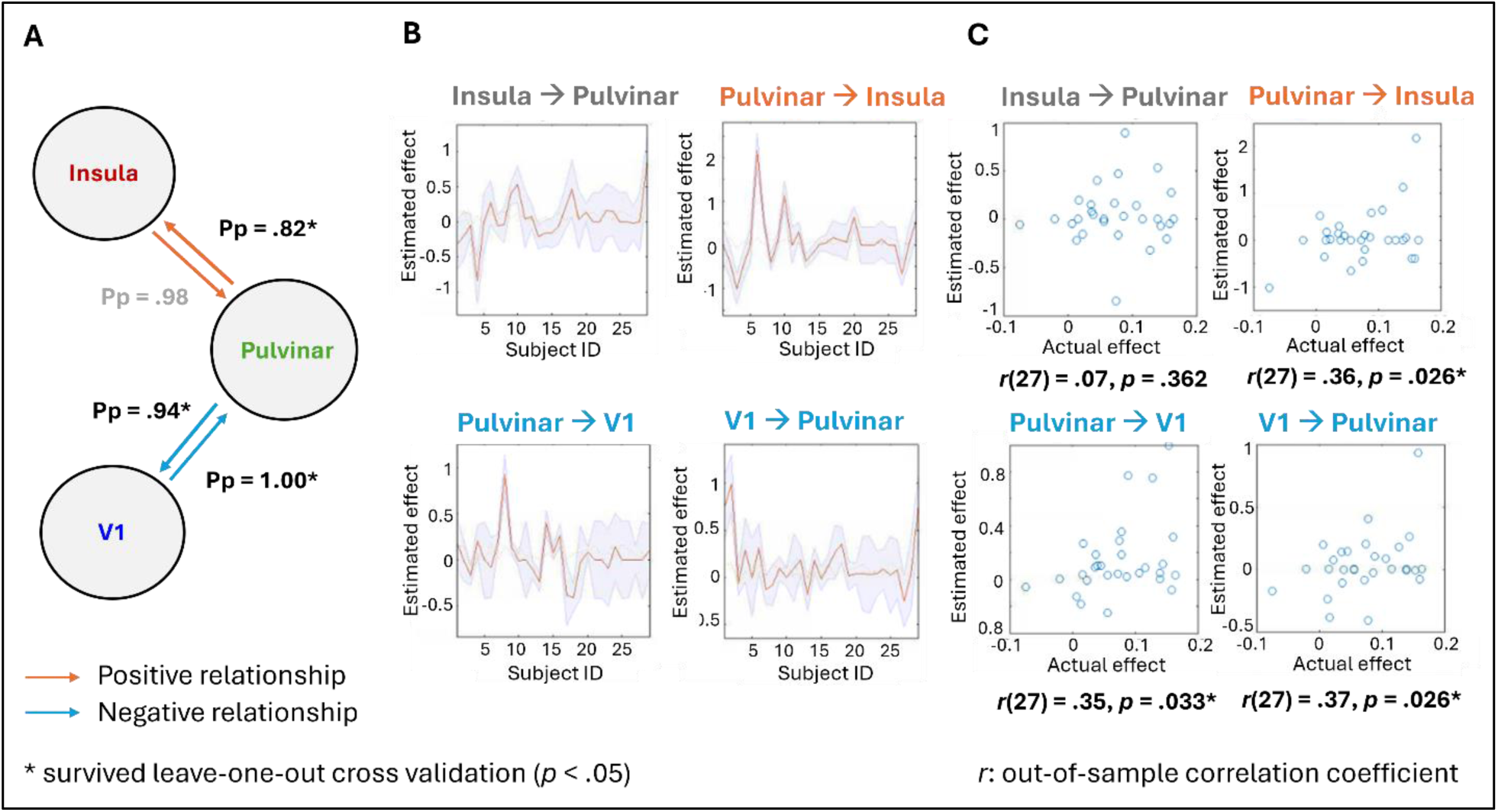
(A) The strength of the modulatory effect of high precision predicted the behavioural effect of high precision (i.e., the difference between participant’s response accuracy in high precision condition and their average response accuracy). Arrows represent relationships with > .75 posterior probability (Pp; the probability that the parameter is different from zero). The grey colour indicates that the modulation did not pass the leave-one-out cross validation. (B) The estimated and mean-centred high-precision effect for each left-out subject (represented by the red line) in the leave-one-out cross validation. The shaded areas represent the 90% confidence interval around the estimated effect. (C) The out-of-samples correlation of the actual scores of the behavioural covariate against the predicted behavioural covariate for each left-out subject. Each dot represents the estimate of a participant’s behavioural covariate.

The connections between the insula and pulvinar positively predicted the high-precision effect: (1) the strength of the modulatory connection from insula to pulvinar (elicited by high precision) positively predicted the high-precision effect (Pp = .98), indicating that a participant with a stronger modulation connection from insula to pulvinar would also tend to show a larger high-precision effect; (2) the strength of the modulatory connection from pulvinar to insula elicited by high precision positively predicted the high-precision effect (Pp = .82), indicating that a participant with a stronger modulation connection from pulvinar to insula elicited by high precision would also tend to show a larger high-precision effect.

The connections between V1 and the pulvinar negatively predicted the high-precision effect: (1) the strength of the modulatory connection from pulvinar to V1 (elicited by high precision) negatively predicted the high-precision effect (Pp = .94), indicating that a participant with a stronger modulation connection from pulvinar to V1 elicited by high precision would tend to show a smaller high-precision effect; (2) the strength of the modulatory connection from V1 to pulvinar negatively predicted the high-precision effect (Pp = 1.00), indicating that a participant with a stronger modulation connection from V1 to pulvinar elicited by high precision would tend to show a smaller high-precision effect.

Leave-one-out cross validation (Figure 7C) further confirmed the predictive validity of three out of the four suprathreshold connections in the BMA results. It was validated that an increase in the strength of the modulatory connection from the pulvinar to insula elicited by high precision would predict an increase in the high-precision effect (*p* = .026). The opposite trend was found in the modulatory connections between the pulvinar and V1. An increase in the strength of the modulatory connection from the pulvinar to V1 elicited by high precision would predict a decrease in the high-precision effect (*p* = .033). An increase in the strength of the modulatory connection from the V1 to pulvinar elicited by high precision would predict a decrease in the high-precision effect (*p* = .026).

## Discussion

This study investigated the neural mechanisms underlying precision modulation in predictive processing during a visual prediction task with varying levels of precision using DCM and 7T fMRI. Consistent with our behavioural hypothesis, participants’ response accuracy was significantly higher in the high-precision condition than in the low-precision condition. In line with our connectivity hypotheses, DCM and PEB analysis showed that high-precision predictions elicited: (1) a modulatory connection from the insula to V1, and (2) from the pulvinar to V1. As hypothesised, high-precision modulation from the pulvinar to V1 was *inhibitory*. However, contrary to our hypothesis, modulation from the insula to V1 was *excitatory*. In addition to the hypothesised modulations, we observed an inhibitory modulation from the insula to the pulvinar. Furthermore, the strength of these modulatory connections could be used to predict the high-precision effect (i.e., the difference between participants’ response accuracy during high-precision predictions and their average response accuracy), highlighting a critical link between effective neural connectivity and behavioural performance.

### The task effectively elicited behavioural differences between precision conditions

Behavioural results revealed significantly higher response accuracy in the high-precision condition than the low-precision condition across participants. This finding indicates that the manipulation of prediction precision within the cueing paradigm effectively induced behavioural differences between precision levels. The observed performance pattern suggests that participants successfully learned the cues and used this information to guide their responses. These results align with the predictive processing framework as higher-precision predictions exert a greater influence on behaviour by being weighted more strongly in perceptual inference (Bastos et al., 2012; Friston, 2010).

### A dual-route feedback mechanism involving insula, pulvinar, and V1

Overall, our findings indicate that high precision of predictions predominantly recruit a dual-route feedback mechanism involving an indirect inhibitory pathway from the insula to V1 via the pulvinar, and a direct excitatory pathway from the insula to V1. The insula has been shown to evaluate the relevance of competing information sources and to prioritise salient information (Huang et al., 2021). In this context, and regarding the inhibitory pathway, the insula may serve to prioritise precise predictions over uncertain sensory input by inhibiting pulvinar activity. Given that the pulvinar has been evidenced to act as a ‘modulator’ that gates V1’s feedforward signalling to higher cortical regions (Purushothaman et al., 2012), inhibition of the pulvinar by the insula may, in turn, reduce V1’s feedforward signalling of the uncertain bottom-up input. In summary, this indirect inhibitory pathway from the insula to V1 via the pulvinar may serve to prioritise and guide top-down predictive processing when the predictions are highly precise.

The direct excitatory pathway from the insula to V1 during high-precision prediction was unexpected, as it diverges from previous findings reporting inhibitory influences from the insula to V1 in response to expected stimuli (Salomon et al., 2016). That said, this excitatory modulation may function to enhance predictive feedback signals to V1, suggesting that the insula plays a dual role, dampening feedforward sensory input via the pulvinar while simultaneously amplifying top-down predictions directly to V1. This dual mechanism may serve to fine-tune the perceptual system by selectively enhancing precise predictions and suppressing imprecise sensory inputs, optimising the balance between prior expectations and incoming data, which is consistent with Bayesian inference models of the brain (Lee & Mumford, 2003; Yon & Frith, 2021).

This dual profile speaks to an ongoing debate in predictive coding frameworks regarding whether predictive feedback leads to sharpening or dampening of sensory representations (Kok et al., 2012). The sharpening account proposes that top-down predictions enhance the representation of expected stimuli by increasing the gain of neurons coding prediction-congruent features in early sensory regions (Kok et al., 2012; Lee & Mumford, 2003), whereas the dampening account suggests that expected input elicits inhibitory influences on early sensory regions (Alink et al., 2010). Our findings may help reconcile this dichotomy by providing evidence for both, as excitatory feedback may sharpen the representation of precise predictions, while inhibitory modulation via the pulvinar may dampen the propagation of imprecise or noisy sensory input. Such a balanced mechanism could be critical for perceptual efficiency and behavioural flexibility, particularly under conditions of high prediction precision. Nonetheless, an important caveat is that the direction of the sign (positive or negative) of DCM parameters reflects relative change in effective connectivity and should not be taken as direct evidence of synaptic excitation or inhibition (Stephan et al., 2010).

Although this study was conducted with neurotypical individuals, the findings may offer insights for understanding aberrant perceptual experiences in neuropsychiatric conditions, such as schizophrenia. Theoretical models of schizophrenia propose that hallucinations and delusions may arise from dysregulated precision weighting in predictive processing, where overly precise priors or unreliable sensory signals lead to maladaptive perceptual inferences (Adams et al., 2013; Corlett et al., 2009; Fletcher & Frith, 2009; Sterzer et al., 2018, 2019). Notably, the opposite pattern has also been reported, with overly imprecise priors leading to underweighted prior beliefs in psychosis (see Goodwin et al. (2025) for a recent review on this topic). Our identification of a dual-route feedback mechanism, whereby the insula enhances top-down predictions and suppresses feedforward sensory input via the pulvinar, offers a potential pathway through which such dysregulations might occur. If, for example, excitatory modulation from the insula to V1 becomes pathologically amplified, or if inhibitory gating via the pulvinar is overactive, the resulting imbalance may favour internally generated predictions over external input, contributing to psychotic-like experiences (Adams et al., 2013; Sterzer et al., 2019). While speculative, these interpretations provide a starting point for future research examining how precision-related connectivity in these regions may be altered in clinical populations.

### Behavioural precision adjustments are driven by excitatory pulvinar to insula inputs and pulvinar-V1 attenuation loops

Our findings reveal relationships between neural dynamics elicited by high precision and their functional relevance at the behavioural level. Specifically, the modulation from the pulvinar to the insula showed a positive relationship with the high-precision effect (i.e., the difference between participants’ response accuracy during high-precision predictions and their average response accuracy). Stronger modulation from pulvinar to insula facilitated performance accuracy during the high-precision condition. This suggests that the adaptive cognitive adjustments associated with high-precision predictions are supported by increased communication from the pulvinar to the insula.

In contrast, the interaction between the pulvinar and V1 demonstrated bilateral negative relationships with the high-precision effect. This indicates that attenuation of the reciprocal connections between the pulvinar and V1 would enhance performance during high-precision conditions. As the connectivity parameters were negative (i.e., inhibitory), a reduction would indicate stronger inhibition. This finding aligns with the proposed theoretical perspective that successful attenuation of V1 feedforward signalling is essential for optimal Bayesian integration when prior predictions are precise and sensory input is uncertain. Conversely, weaker thalamocortical inhibition could reduce the behavioural benefit under high-precision conditions, indicating insufficient prioritisation of prior predictions over sensory input. These insights further supported the observed effective modulations elicited by high-precision predictions.

### Limitations and future directions

This study has several limitations. One limitation lies in the use of DCM itself, which is constrained by computational and interpretational limits on the number of brain regions that can be included in the model (Lohmann et al., 2012). Notably, regions such as the inferior frontal gyrus (IFG), which are implicated in top-down feedback processing (Ficco et al., 2021), were not included in the model space. The secondary visual cortex (V2), which has been proposed to function both as a relay and as a computational hub for comparing and refining bottom-up sensory inputs from V1 with top-down predictions from higher-order visual areas (Shipp, 2023) was also excluded. As this study served as a starting point to explore precision modulation across brain regions, we opted to confine our analysis to a focused, theory-driven subset of regions. Future research could expand upon this work by incorporating additional regions to more comprehensively understand precision modulation in the brain’s predictive processing network.

Precision modulation of *prediction errors* is another key mechanism for learning about the contingencies in our environment and updating our internal model (Friston, 2005). In this study, however, we focused on precision modulation for *predictions* specifically, thus designing the task as to maximise the number of trials and hence the statistical power of our analyses. To that end, participants were shown feedback about their performance only once every ten trials rather than after each trial. However, to investigate the precision modulation for prediction errors, we would need to present feedback to participants immediately after each trial, thus probing a prediction error response when the response was incorrect. Future studies could incorporate trial-by-trial feedback and increased temporal spacing around the feedback window to investigate the neural mechanisms of precision modulation in the context of prediction errors and gain a deeper understanding of precision in visual predictive processing.

## Conclusion

In summary, this study found that precision modulation in visual predictive processing is driven by modulatory connectivity between the insula, pulvinar, and V1. High-precision predictions elicited excitatory modulation from the insula to V1 and inhibitory modulation from the insula to the pulvinar and from the pulvinar to V1. We proposed a dual-route feedback mechanism whereby the insula enhances top-down feedback to V1 while indirectly suppressing bottom-up feedforward signalling in V1 via the pulvinar. This mechanism may serve to optimise the balance between prior predictions and incoming sensory input, aligning with Bayesian inference models of perception (Lee & Mumford, 2003; Yon & Frith, 2021).

Critically, this network of modulatory connections demonstrated behavioural relevance. During high-precision predictions, a stronger connection from pulvinar to insula and weaker bilateral connections between the pulvinar and V1 predicted adaptive cognitive adjustments. This suggests that enhanced pulvinar-insula communication, alongside effective attenuation of V1 feedforward signalling, may be critical for optimal Bayesian integration when prior predictions are precise and sensory input is uncertain. Together, these results underscore the importance of the modulatory connectivity between the insula, pulvinar, and V1 in the precision modulation of visual predictions, advancing our understanding of the neural implementation of predictive processing and offering potential implications for investigating atypical precision-weighting mechanisms in clinical populations.

## Data availability statement

The data that support the findings of this study are openly available in GitHub at https://github.com/linzhi-tao/Precision_study.

## Ethics approval statement

This study was approved by the University of Melbourne Human Research Ethics Committee (Ethics ID: 24710, reference number: 2022-24710-35008-4). All participants provided written informed consent and received monetary compensation for participation.

## Acknowledgment

The authors acknowledge the facilities, scientific and technical assistance from the National Imaging Facility, a National Collaborative Research Infrastructure Strategy capability, at the Melbourne Brain Centre Imaging Unit, The University of Melbourne. This work was supported by a research collaboration agreement with Siemens Healthineers.

