## Supporting Information for "Effective connectivity reveals dual-route mechanism of visual prediction precision via insula and pulvinar"

| Peak MNI Coordinates (mm) | | | Cluster Size (voxels) | Cluster *p* (uncorrected) | Peak *T* | Anatomic Region Label |
|---|---|---|---|---|---|---|
| *x* | *y* | *z* | | | | |
| ***High precision > Low precision*** | | | | | | |
| 10 | -82 | 5 | 9938 | 0 | 5.59 | Calcarine R |
| 10 | -72 | -8 | | | 4.8 | Lingual R |
| 6 | -90 | 16 | | | 4.57 | Cuneus R |
| -53 | 0 | 5 | 241 | 0.012 | 3.82 | Rolandic Oper L |
| -59 | 8 | -2 | | | 2.97 | Temporal Pole Sup L |
| 2 | -59 | 77 | 49 | 0.217 | 3.8 | Precuneus R |
| 10 | -43 | 53 | 1925 | 0 | 3.78 | Precuneus R |
| 43 | -26 | 50 | | | 3.6 | Postcentral R |
| 8 | -26 | 70 | | | 3.24 | Paracentral Lobule R |
| -16 | 30 | 64 | 40 | 0.264 | 3.68 | Frontal Sup 2 L |
| -59 | -5 | 11 | 127 | 0.056 | 3.63 | Rolandic Oper L |
| -51 | -18 | 14 | | | 2.23 | Rolandic Oper L |
| -53 | -10 | 50 | 1984 | 0 | 3.5 | Postcentral L |
| -27 | -61 | 62 | | | 3.3 | Parietal Sup L |
| -27 | -54 | 54 | | | 3.25 | Parietal Inf L |
| -34 | -3 | 3 | 52 | 0.204 | 3.5 | Insula L |
| -14 | -14 | 24 | 15 | 0.5 | 3.4 | Caudate L |
| -51 | -30 | -21 | 11 | 0.569 | 3.38 | Temporal Inf L |
| 19 | -19 | 5 | 68 | 0.15 | 3.28 | Thal VPL R |
| 30 | -26 | 6 | | | 2.75 | Insula R |
| ***Low precision > High precision*** | | | | | | |
| -26 | -29 | 40 | 127 | 0.056 | 3.71 | Postcentral L |
| -16 | -22 | 43 | | | 3.7 | Cingulate Mid L |
| -8 | -22 | 37 | | | 2.18 | Cingulate Mid L |

| | | | | | | |
|---|---|---|---|---|---|---|
| 46 | -70 | -14 | 22 | 0.409 | 3.61 | Occipital Inf R |
| -22 | 8 | -21 | 116 | 0.066 | 3.57 | OFCpost L |
| -24 | -3 | -21 | | | 2.84 | Amygdala L |
| -18 | 3 | -14 | | | 2.48 | Olfactory L |
| 10 | 16 | 8 | 168 | 0.031 | 3.5 | Caudate R |
| 3 | 3 | 16 | | | 2.78 | Caudate R |
| 0 | 10 | 10 | | | 2.18 | Caudate L |
| -29 | -32 | -8 | 405 | 0.002 | 3.46 | Hippocampus L |
| -24 | -51 | -6 | | | 2.99 | Lingual L |
| -35 | -43 | -10 | | | 2.74 | Fusiform L |
| -30 | -10 | 32 | 26 | 0.368 | 3.42 | Precentral L |
| 32 | -19 | -8 | 24 | 0.387 | 3.25 | Hippocampus R |

**Table S1.** Peak activations of the second-level differences between high- and low-precision conditions. Activations were thresholded at $p < .05$, pTFCE-corrected (Spisák et al., 2019), with a minimum cluster size (k) = 10. Anatomic region labels were obtained via Automated Anatomical Labelling Atlas 3 (AAL3; Rolls et al., 2020). MNI: Montreal Neurological Institute. Sup: superior. Oper: operculum. Inf: inferior. Mid: middle. L: left. R: right. Thal: thalamus. VPL: ventral posterolateral. OFC: orbitofrontal cortex. post: posterior.
